# Baseline human gut microbiota profile in healthy people and standard reporting template

**DOI:** 10.1101/445353

**Authors:** Charles Hadley King, Hiral Desai, Allison C. Sylvetsky, Jonathan LoTempio, Shant Ayanyan, Jill Carrie, Keith A. Crandall, Brian C. Fochtman, Lusine Gasparyan, Naila Gulzar, Paul Howell, Najy Issa, Konstantinos Krampis, Lopa Mishra, Hiroki Morizono, Joseph R. Pisegna, Shuyun Rao, Yao Ren, Vahan Simonyan, Krista Smith, Sharanjit VedBrat, Michael D. Yao, Raja Mazumder

## Abstract

A comprehensive knowledge of the types and ratios of microbes that inhabit the healthy human gut is necessary before any kind of pre-clinical or clinical study can be performed that attempts to alter the microbiome to treat a condition or improve therapy outcome. To address this need we present an innovative scalable comprehensive analysis workflow, a healthy human reference microbiome list and abundance profile (GutFeelingKB), and a novel Fecal Biome Population Report (FecalBiome) with clinical applicability. GutFeelingKB provides a list of 157 organisms (8 phyla, 18 classes, 23 orders, 38 families, 59 genera and 109 species) that forms the baseline biome and therefore can be used as healthy controls for studies related to dysbiosis. The incorporation of microbiome science into routine clinical practice necessitates a standard report for comparison of an individual’s microbiome to the growing knowledgebase of “normal” microbiome data. The FecalBiome and the underlying technology of GutFeelingKB address this need. The knowledgebase can be useful to regulatory agencies for the assessment of fecal transplant and other microbiome products, as it contains a list of organisms from healthy individuals. In addition to the list of organisms and abundances the study also generated a list of contigs of metagenomics dark matter. In this study, metagenomic dark matter represents sequences that cannot be mapped to any known sequence but can be assembled into contigs of 10,000 nucleotides or higher. These sequences can be used to create primers to study potential novel organisms. All data is freely available from https://hive.biochemistry.gwu.edu/gfkb and NCBI’s Short Read Archive.

## INTRODUCTION

While humanity has only begun to influence planetary-level events in the last few hundred years [1], microorganisms have shaped our planet since time immemorial [2]. It has been shown that the microbes of the ocean are as important for influencing planetary climate as the microbes of gastrointestinal (GI) tracts of cattle [3]; furthermore, new functions are continuously found for the human microbiome [4-6]. However, since the advent of germ theory and the antimicrobial chemotherapy revolution, microbes have been viewed as insurgents bound for eradication [7].

In 2001, some sixty years into the antibiotic era, Joshua Lederberg coined the term ‘microbiome’ as the pendulum of opinion began to swing back to a more microbe-tolerant position [8,9]. In 2008, the US National Institutes of Health launched the Human Microbiome Project (HMP) to better understand the makeup of the community of microbes in cohabitation with humans [10,11]. This population of microorganisms brings with it a vast, diverse, and modifiable set of genomes which have proven to influence human health and disease [12,13]. Together, these organisms’ genomes comprise the metagenome, a highly versatile pool of genetic elements which now serves as a target for medical research [14]. Microbiome characterization through various analysis pipelines has advanced progressively since HMP and this development process has catalyzed the understanding of certain roles of these microbial communities [15,16].

Although, microbiomes of all body sites are important, the gut microbiome with hundreds of prevalent species is of major interest to a large and diverse number of researchers [17,18]. The healthy gut microbiome data and analysis is crucial for all studies of disease with relation to the human gut. A *Nature Microbiology* issue in 2016 contained a consensus statement which outlined all federally-funded microbiome research over a three-year period [19]. The authors, on behalf of the federal government’s FastTrack Action Committee on Mapping Microbiomes (FTAC-MM), defined a microbiome as a multispecies community of microorganisms in any environment: host, habitat, or ecosystem. One of the conclusions reached by the authors was a “priority need” for higher-throughput, more accurate data acquisition, better pipelines for data analyses, and a greater ability to organize, store, access, and share/integrate data sets. At present, most studies leverage study specific control groups and reporting mechanisms. This problem is compounded by the fact that different bioinformatics pipelines produce different results largely because all current pipelines use a limited number of *ad hoc* reference organisms to determine abundance. It has also been shown that database growth influences the accuracy of relatively faster k-mer-based species identification [20]. The final understanding of the baseline healthy microbiome therefore can be flawed because the methods are uniquely applied in each study. As such, there is a need for aggregation, validation for interoperability, and eventual standardization of methods and reporting.

Currently, all metagenomic analyses use as a reference database, nucleotide sequences from a limited set of pre-determined microorganisms or genes and, as such, these reference lists are not truly comprehensive. The use of limited sets of sequence data is prevalent because it is computationally challenging to perform pairwise read alignment against the entire NCBI non-redundant nucleotide database (NCBI-nt) [21]. We have developed algorithms that allow the use of the complete NCBI-nt and have shown that using the NCBI-nt allows accurate analysis of the data with significantly less errors in microorganism abundance quantification [22]. To leverage this prior work on metagenomics analysis algorithm, we collected and sequenced healthy cohort of samples from participants. To make sure the samples are abundant and correct enough to build healthy reference list, we also retrieved sequences of healthy people from HMP. The method also generates a list of assembled contigs that cannot be aligned to any known sequence in NCBI-nt but are present in healthy individual fecal samples and are ideal for healthy-disease-microbiome correlation analysis and novel primer design. We define these sequences as metagenomic dark matter – sequences that cannot be mapped to any known sequence but can be assembled into contigs of 10,000 nucleotides or higher. The contig nucleotide length threshold is expected to reduce the number of contigs in GutFeelingKB that are not of biological origin. Our definition is much stricter than previous definitions of the metagenomic dark matter which accepts remote homology to known sequences [23]. The need to include metagenomic dark matter in comprehensive analyses of the gut microbiome matches the arguments presented by Bernard et al. in their recent manuscript on microbial dark matter where they opine that “unraveling the microbial dark matter should be identified as a central priority for biologists” [24].

Together, our methods and GutFeelingKB with significant new data, allow for the analysis of the species-level composition of the healthy human gut microbiome and also the metagenomic dark matter. We have subsequently designed a standard reporting template of individual microbiome data to be compared to the database, useful to any scientist, clinician, or patient.

## MATERIALS AND METHODS

### Metagenomic sampling and participant statistics

#### Healthy cohort selection and nutritional information

Participants for this study were recruited from the George Washington University (GW) Foggy Bottom campus area through the use of flyers and emails to GW affiliated organizations (selection criterions included in S1 Table). Study participants provided samples and anthropomorphic measurements (included in S1 Table) were collected from healthy people at the George Washington University according to an IRB approved protocol. At the baseline visit, participants received extensive instructions on how to record their dietary intake (including type, brand, and portion size of every food and beverage consumed on each day throughout the study period) and the time of consumption for each item. Participants then recorded their dietary intake using a seven-day food journal throughout the study. Each participant provided three samples. The food journal was collected at the submission of the final sample, after which the reported 7-day dietary intakes for each subject were entered into the Nutrition Data System for Research (NDSR) [25]. NDSR produces a tabular daily nutrient for each day of dietary intake for each individual, which was then added as metadata to the abundance matrices (supplementary table S2 Table). All participants self-reported as ‘healthy’ at the start of the study and remained healthy throughout.

#### Sampling and sequencing

Fecal samples were collected from healthy volunteers using sterile commode containers at the Milken Institute School of Public Health at the George Washington University (GWSPH). Immediately following collection, fecal samples were stored in a −20 degree Celsius freezer for a period of up to two weeks, after which, aliquots were placed in longer term storage at −80 degree Celsius ultra-freezer. Samples were subsequently transported to the sequencing center on dry ice. DNA was extracted using the MoBio PowerFecal DNA Isolation kit25. Double-stranded DNA (dsDNA) concentration and quality was assessed using NanoDrop and the Qubit dsDNA Broad Range (BR) DNA Assay Kit26, respectively. DNA was diluted for library preparation using the Illumina Nextera XT Library Prep Kit, and 1 ng from each sample was fragmented and amplified using Illumina Nextera XT Index Kit primers. Amplified DNA was then cleaned using Agencourt AMPure XP beads, resuspended in buffer, and tested again for concentration, quality, and fragment size distribution on a Bioanalyzer using the Agilent High Sensitivity DNA Kit. DNA libraries were brought to the same nM concentration, pooled, and denatured with 0.2 N NaOH prior to loading on an Illumina MiSeq Reagent Kit v3 and sequencing on the Illumina MiSeq platform. Sequence data FASTQ files was uploaded to BaseSpace (https://basespace.illumina.com/home/index) for sharing and further analysis.

#### Sequence quality assurance

All sequence data were uploaded to the GW High-performance Integrated Virtual Environment (HIVE) [26,27]. Upon initial upload into the system, HIVE conducts a series of quality assurance (QA) computations for each sequence read file and generates figures to display the results. S3 Fig shows the quality assurance computations done on one read file.

Upon completion of the initial loading, quality analysis resulting figures were inspected for each read file to ensure that the read file was of adequate quality and did not have any unusual characteristics such as low quality score or disproportionate ratio and distribution of A, T, G and C nucleotides. S4 Fig shows the aggregated computations across all samples.

#### Healthy cohort from Human Microbiome Project

In addition to the data generated from sequencing described above, additional data were downloaded and analyzed from the Human Microbiome Project (HMP) [28]. HMP sequence data and metadata are available through NCBI SRA and dbGaP. We obtained fifty fecal metagenomic samples, randomly chosen from HMP Phase I (supplementary table S1 Table) to match approximately the number of samples collected in our study. For the samples collected by us and the HMP project dataset subjects were screened based on stringent criteria; the individuals who passed screening were considered “healthy” subjects[11].

#### GW and HMP combined data

Sequence and metadata from this study are publicly available through GutFeeling (https://hive.biochemistry.gwu.edu/gfkb), and also available from NCBI-SRA BioProject Healthy Human Gut Metagenomics (PRJNA428202), and Effects of non-nutritive sweeteners on the composition of the human gut microbiome (PRJNA487305). For PRJNA487305, samples prior to intake of non-nutritive sweeteners were used in this study. HMP data was downloaded from NIH Human Microbiome Project (HMP) Roadmap Project (PRJNA43021).

48 samples from 16 individuals were sequenced in the GW cohort. Each sample resulted in two pair-end read files (for details see S5 Table). Sequence data from these 48 samples along with 50 samples from HMP passed sequence quality checks and was used to develop the baseline microbiota profile. For GW samples 55.55% (± 13.46%) while for HMP 48.29% (± 18.54%) of the reads could not be mapped to any known sequence. There was no need for any computational filtering of human DNA as the MoBio PowerFecal DNA Isolation kit25 was used for GW samples, biochemically removing any host DNA. The human DNA had been computationally removed before the HMP data was deposited in dbGaP[11].

Data interoperability is a perennial challenge in bioinformatics [29]. This problem is further magnified when considerations are made for data from samples collected in distant locations at different times. In the case of HMP, sampling was done in Houston, TX and St. Louis, MO during 2008-2012. All GW samples were collected from the DC Metro Area in 2016. One way to test the compatibility of these data sets was to run a Between-Class Analysis (BCA) on all samples from each of the projects. Data from our three, separate projects fell into the expected three classic enterotypes [30] instead of clustering by project set (S6 Fig). Had the data clustered by project, sampling location, or year, they may not have been compatible for inclusion in the same database. However, we believe that these data do not show a sampling bias and can be leveraged for joint analysis. Sample and participant information can be seen in Table 1.

**Table 1:** Human Microbiome Project (HMP) and GW participant statistics.

| Feature | White | Other | Asian | Black | Male | Female |
| --- | --- | --- | --- | --- | --- | --- |
| HMP samples | 39 | 2 | 7 | 2 | 30 | 20 |
| GW samples | 24 | 0 | 6 | 18 | 21 | 27 |

### Filtered-nt

The Filtered-nt (v3.6) was created from the NCBI-nt file through the use of taxonomy blacklist file. NCBI-nt and NCBI taxonomy files were downloaded (ftp://ftp.ncbi.nlm.nih.gov/blast/db/FASTA/; ftp://ftp.ncbi.nlm.nih.gov/pub/taxonomy) on May 21st, 2017. More specifically, Filtered-nt was generated using blacklist file of taxonomy IDs identified based on terms that are contained in the lineage of each taxonomy entry. Taxonomy nodes with terms such as ‘unclassified’, ‘unidentified’, ‘uncultured’, ‘unspecified’, ‘unknown’, ‘vector’, ‘environmental sample’, ‘artificial sequence’, ‘other sequence’ were blacklisted. Child nodes are also automatically blacklisted. The filtered taxonomy list was then used to filter the NCBI-nt sequence file. Filtered-nt and the blacklisted taxonomy IDs along with node names are available for download at hive.biochemistry.gwu.edu/filterednt.

### Metagenomic analysis pipeline

The innovative metagenomic analysis pipeline we developed includes three software tools and one sequence database (Filtered-nt), organized in a fashion to produce a workflow that ensure an efficient and comprehensive analysis of a large sequence space. The tools are CensuScope [22], HIVE-Hexagon [31], and IDBA-UD [32]. All software tools are integrated in the HIVE platform [26,27] and allow end-to-end analysis of metagenomic sequences.

#### Bacterial abundance profile

Figure 1 provides a schematic representation of the workflow. The first step uses CensuScope to identify organisms that are present in the sample [22]. CensuScope is a taxonomic profiling software that randomly extracts a user defined number of reads and maps them to any size sequence database using BLAST [33]. In our previous studies, we have shown that CensuScope is rapid, accurate and is not hindered by the size of the reference sequence database. With the non-redundant sequence database’s almost constant exponential increase, CensuScope offers a scalable approach for estimating taxonomic composition of a microbial population. We then used HIVE-hexagon, a highly specific and sensitive short-read aligner [31], to obtain the final abundance profiles. HIVE-hexagon was used to map all the reads to the organisms that are identified through CensuScope.

**Figure 1.**
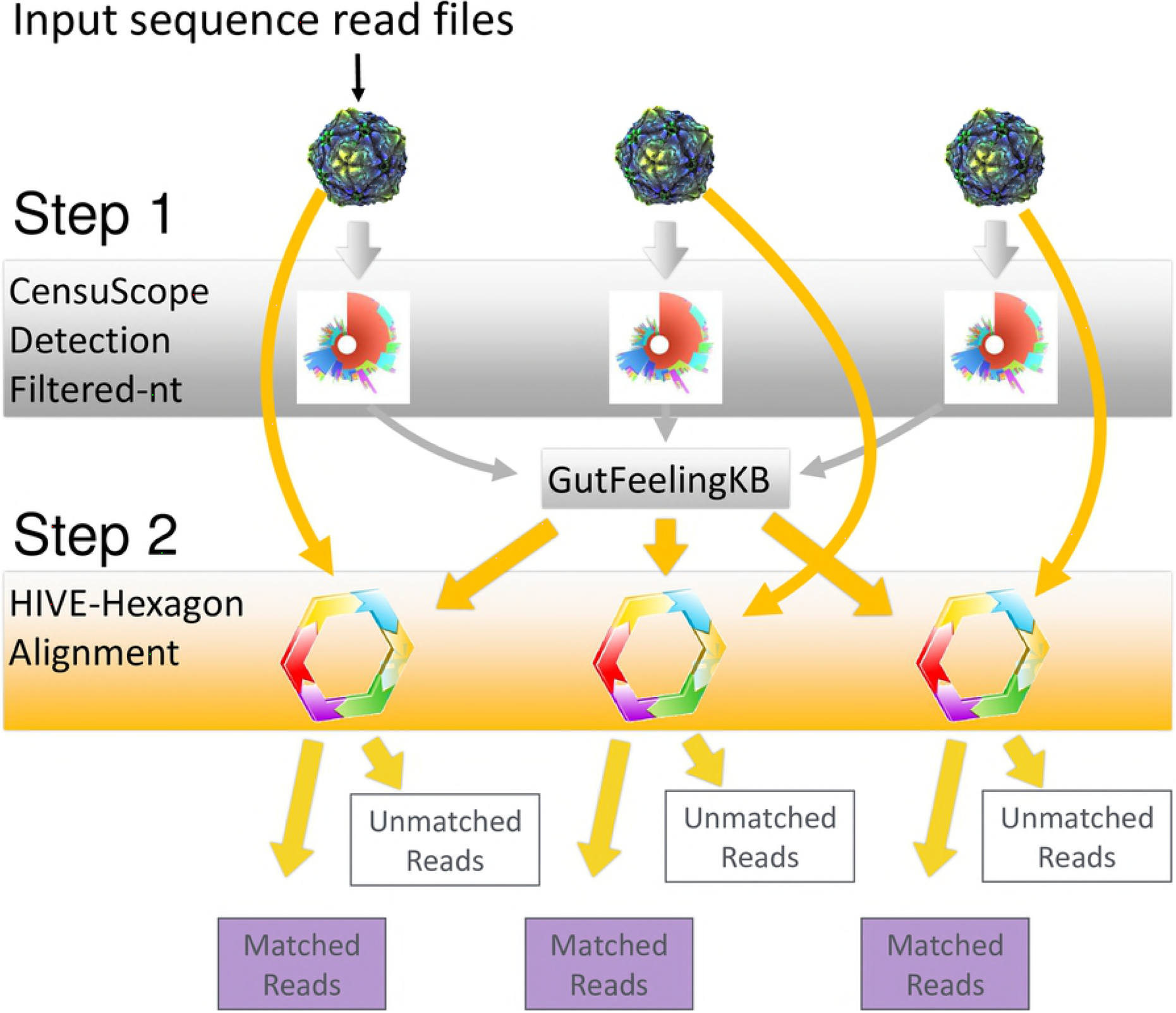
Metagenomic analysis pipeline. Step 1: CensuScope is run for each read file against Filtered-nt. Each of the aligned organism approved by manually check will be added to the GutFeelingKB and it is versioned. Step 2: For the final analysis the raw read files are run in HIVE-hexagon against the GutFeelingKB and the outputs are tabulated as relative abundance percentages.

#### Healthy Human gut microbiome list (GutFeelingKB)

A list of organisms and taxonomy identifiers are provided as the output by CensuScope. After manual verification that the alignment results are valid for each of the identified organisms, every new organism and their alignments are checked manually to confirm that it is a true positive. Manual evaluation includes match count (number of matched alignments over the entire computation (all iterations)), valid taxonomy level assignment, completeness of sequence and contamination in genome assembly in Filtered-nt. The accession numbers are then used to obtain the NCBI Genome Assembly IDs, which is used to retrieve proteome IDs from UniProt whenever possible. Genome to proteome mapping was guided by Representative Proteome Groups (RPGs), a dataset that contains similar proteomes (hence genomes) calculated based on co-membership in UniRef50 clusters [34] (supplementary table S7 Table). Such mapping provides an opportunity to explore metabolic pathways present in the identified organisms. It is important to note that many bacteria are closely related and hence have large homologous regions. This can lead to species level misidentification. Although the concept of pan-genome or pan-proteome for closely related bacteria is well accepted [35], it is important to avoid such misidentification for known pathogens. To avoid such false positives of well-known pathogens (S8 Table), they are included only if their abundance is 1% or higher and their alignments have been manually checked.

#### Metagenomic dark matter

The unaligned reads of each sample were assembled using IDBA-UD [32]. Assembled contigs longer than 10,000 nucleotides were considered as metagenomic dark matter. Such a large length threshold was used to ensure that the metagenomics dark matter contigs are of biological origin. The gut microbiome of a sample can be represented as the sum of known organisms and organisms represented by the metagenomic dark matter sequences. More specifically, the contigs that were over 10,000 nucleotides in length were tagged with the sample ID and numbered, and metadata data about the participant were added to the header. These contigs are available as a download at (https://hive.biochemistry.gwu.edu/gfkb) for further analysis and novel primer design.

### Analysis of nutritional metadata and microbial abundance

MaAsLin, an R package that employs a “multivariate statistical framework that finds associations between clinical metadata and microbial community abundance or function” [36] was used to find correlations between bacterial abundance and diet. Intra-host variability was analyzed evaluating the standard deviation of multiple measurements for every patient averaged over all patients. Inter-host variability was computed as a standard deviation of the means of per-host abundance values. To estimate the degree of stability of measurements for bacterial populations in patient samples intra-host vs inter-host variability ratio was computed.

Nutrition to organism abundance correlation was also computed by using a Cosine Similarity Coefficient. The matrix of bacterial strain abundances was variance scaled and zero centered to create comparable distributions of equal variability. Categorical data (such as gender) was turned into numerical values. More specifically, in order to define correlation metrics between features and bacterial composition for the set of individuals, we used Cosine Similarity Coefficient as defined in Formula 1.

#### Formula 1

Cosine Similarity Coefficient of correlation between bacteria j and feature k is computed as the sum product of j-th Bacteria (Bj) abundance for patient i and k-th Feature (Fk) of patient i.

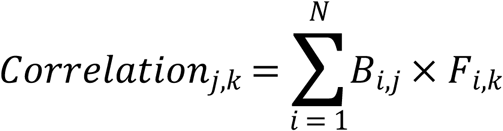

A Cosine Similarity of around 1 means strong correlation, −1 means strong anti-correlation, 0 means no correlation with 0.7 being considered the marginal threshold for evidence of some degree of correlation.

## RESULTS AND DISCUSSIONS

### Filtered NCBI-nt (Filtered-nt)

NCBI nucleotide sequence collection (NCBI-nt) is the most comprehensive collection of DNA sequences [21], but many sequences present in NCBI-nt do not provide enough relevant information or they might be artificial (e.g. sequences with taxonomy placement such as environmental, unclassified, synthetic sequences, unidentified sequences etc.). Reads mapped to such sequences do not provide any valuable information in terms of the organisms and hence are not useful in understanding the microbial composition of the sample. The NCBI-nt initially contained 42,439,338 sequences. The taxonomy file contained 1,601,859 scientific names. After removal of 250,610 blacklisted taxonomy IDs (supplementary table S9 Table) containing 7,499,592 sequences the Filtered-nt contained 34,939,806 sequences. The Filtered-nt is ideal for comprehensive metagenomic analysis that relies on best sequence hit.

All current studies use genomes from known gut bacteria as reference database [18,22,37,38] and hence would not be able to detect organisms that are not present in the reference database. The use of Filtered-nt guarantees that no known organism in the sample is missed.

### Healthy fecal microbiome

#### GutFeelingKB - a reference list for healthy human gut organisms

GutFeelingKB consists of 157 organisms which fall into sixty distinct genera, as seen in Table 2 which list in species level and the full table that can be downloaded at https://hive.biochemistry.gwu.edu/gfkb. Members of the Firmicutes and Bacteroidetes phyla make up a majority of the bacterial species were present in the human intestinal microbiota. A total of 155 bacterial and 2 archaeal species were identified in healthy samples. In summary, the healthy human gut microbiome consists of 8 phyla, 18 families, 23 classes, 38 orders, 59 genera and 109 species. 63 (40%), 32 (20%) and 31 (19.7%) members belongs to Firmicutes, Actinobacteria and Bacteroidetes, respectively which make up a majority of the bacterial species. More than half of Firmicutes are members of the Clostridia (20.3%) class, which is the most abundant class, followed by Bacteroidia (18.5%), Bifidobacteriales (16.6%), Enterobacterales (14%) and Lactobacillales (14%). All of members of Clostridia in the samples are members of Clostridiales order and all of Bacteroidia belongs to Bacteroidales, these two are the most abundant orders. There are 27 organisms are members of Bifidobacteriaceae family, and 26 of them belongs to *Bifidobacterium longum*, which are the most abundant species.

**Table 2.** List of 109 baselines species and their GenBank accessions found in healthy human gut.

| Organism name | GenBankAC | Organism name | GenBankAC | Organism name | GenBankAC |
| --- | --- | --- | --- | --- | --- |
| Acidaminococcus fermentans (Bac/Firmicute) (100<sup>1</sup>;0.04<sup>2</sup>) | CP001859 | Clostridium saccharolyticum (Bac/Firmicute) (100;0.24) | CP002109, FP929037 | Odoribacter splanchnicus (Bac/CFB_bac) (100;1.12) | CP002544 |
| Acidaminococcus intestine (Bac/Firmicute) (100;0.09) | CP003058 | Coprococcus catus (Bac/Firmicute) (100;0.37) | FP929038 | Ornithobacterium rhinotracheale (Bac/CFB_bac) (100;0.11) | CP006828 |
| Acidovorax sp KKS102 (Bac/Beta-proteo) (100;0.01) | CP003872 | Coprococcus sp ART55/1 (Bac/Firmicute) (100;0.68) | FP929039 | Oscillibacter valericigenes (Bac/Firmicute) (100;0.05) | AP012044 |
| Adlercreutzia equolifaciens (Bac/ActnBac) (100;0.07) | AP013105 | Cutibacterium acnes (Bac/ActnBac) (100;0.004) | CP003084 | Paenibacillus sabinae (Bac/Firmicute) (100;0.01) | CP004078 |
| Akkermansia muciniphila (Other Bacteria) (91.84;0.70) | CP001071 | Eggerthella lenta (Bac/ActnBac) (100;0.04) | CP001726 | Paeniclostridium sordellii (Bac/Firmicute) (100;0.02) | LN679998, LN681234 |
| Alistipes finegoldii (Alistipes finegoldii) (100;1.27) | CP003274 | Eggerthella sp. YY7918 (Bac/ActnBac) (100;0.01) | AP012211 | Parabacteroides distasonis (Bac/CFB_bac) (100;2.30) | CP000140 |
| Alistipes shahii (Bac/CFB_bac) (100;1.75) | FP929032 | Enterococcus faecium (Bac/Firmicute) (100;0.04) | CP003351, CP006620, CP006030 | Parvimonas micra (Bac/Firmicute) (100;0.01) | CP009761 |
| Anaerococcus prevotii (Bac/Firmicute) (100; 0.003) | CP001708 | Enterococcus hirae (Bac/Firmicute) (96.94;0.004) | CP003504 | Porphyromonas asaccharolytica (Bac/CFB_bac) (98.98;0.01) | CP002689 |
| Anaerostipes hadrus<br>(Bac/Firmicute)<br>(100;0.55) | FP929061 | Escherichia coli<br>(Bac/Gamma-proteo) (100;1.87) | CP009859, CP010816, CP000948, CP001637, CP000970, CP000243, CP009166, CP002291, CP003297, CP007394, AP009378, AE014075, CP010371, CP002729, CP007799, CP001396, CP009789, CP004009, CP007390, FN649414, CP009167, HG941718 | Porphyromonas gingivalis<br>(Bac/CFB_bac) (100;0.01) | AP009380 |
| Bacillus methanolicus<br>(Bac/Firmicute)<br>(100;0.01) | CP007739 | Escherichia coli O104:H4<br>(Bac/Gamma-proteo) (96.94;0.04) | CP004009 | Prevotella dentalis<br>(Bac/CFB_bac) (100;0.08) | CP003368, CP003369 |
| Bacteroides cellulosilyticus<br>(Bac/CFB_bac)<br>(100;3.38) | CP012801 | Escherichia coli O83:H1<br>(Bac/Gamma-proteo) (95.92;0.06) | CU651637 | Prevotella denticola<br>(Bac/CFB_bac) (98.98;0.04) | CP002589 |
| Bacteroides dorei<br>(Bac/CFB_bac)<br>(100;17.44) | CP007619, CP009057 | Ethanoligenens harbinense<br>(Bac/Firmicute)<br>(100;0.01) | CP002400 | Prevotella intermedia<br>(Bac/CFB_bac) (100;0.07) | AP014597, CP003502, CP003503, AP014598 |
| Bacteroides fragilis<br>(Bac/CFB_bac)<br>(100;3.47) | FQ312004, CR626927, AP006841, AP006842, CR626928 | Eubacterium eligens<br>(Bac/Firmicute)<br>(100;0.65) | CP001104, CP001105, CP001106 | Prevotella melaninogenica<br>(Bac/CFB_bac) (100;0.24) | CP002122, CP002123 |
| Bacteroides helcogenes<br>(Bac/CFB_bac)<br>(100;0.50) | CP002352 | Eubacterium limosum<br>(Bac/Firmicute)<br>(100;0.03) | CP002273 | Prevotella ruminicola<br>(Bac/CFB_bac) (100;0.06) | CP002006 |
| Bacteroides ovatus<br>(Bac/CFB_bac)<br>(100;7.72) | CP012938 | [Eubacterium] rectale<br>(Bac/Firmicute)<br>(100;6.21) | FP929042, FP929043, CP001107 | Prevotella sp oral taxon 299<br>(Bac/CFB_bac) (100;0.06) | CP003666 |
| Bacteroides salanitronis<br>(Bac/CFB_bac)<br>(100;0.48) | CP002530 | [Eubacterium] siraeum<br>(Bac/Firmicute)<br>(100;0.75) | FP929044, FP929059, | Raoultella ornithinolytica<br>(Bac/Gamma-proteo) (100;0.01) | CP004142 |
| Bacteroides sp. CAG:98<br>(Bac/CFB_bac)<br>(100;8.89) | CP008741 | Faecalibacterium prausnitzii<br>(Bac/Firmicute)<br>(100;3.52) | FP929045, FP929046 | Roseburia hominis<br>(Bac/Firmicute) (100;0.69) | CP003040 |
| Bacteroides<br>thetaiotaomicron<br>(Bac/CFB_bac)<br>(100;3.78) | AE015928,<br>AY171301 | Faecalitalea<br>cylindroides<br>(Bac/Firmicute)<br>(100;0.15) | FP929041 | Roseburia<br>intestinalis<br>(Bac/Firmicute)<br>(100;1.15) | FP929049,<br>FP929050 |
| Bacteroides<br>vulgatus<br>(Bac/CFB_bac)<br>(100;14.99) | CP000139 | Fermentimonas<br>caenicola<br>(Bac/CFB_bac)<br>(100;0.01) | LN515532 | Rubinisphaera<br>brasiliensis<br>(Bac/Plnctmy)<br>(70.41;0.0002) | CP002546 |
| Bacteroides<br>xylanisolvens<br>(Bac/CFB_bac)<br>(100;4.92) | FP929033 | Gardnerella<br>vaginalis<br>(Bac/ActnBac)<br>(91.84;0.002) | CP001849 | Ruminococcus<br>bicirculans<br>(Bac/Firmicute)<br>(100;2.54) | HF545616,<br>HF545617 |
| Barnesiella<br>viscericola<br>(Bac/CFB_bac)<br>(100;0.33) | CP007034 | Gordonibacter<br>pamelaeae<br>(Bac/ActnBac)<br>(100;0.03) | FP929047 | Ruminococcus<br>bromii<br>(Bac/Firmicute)<br>(100;0.83) | FP929051 |
| Bifidobacterium<br>adolescentis<br>(Bac/ActnBac)<br>(97.96;0.46) | CP007443,<br>CP010437,<br>AP009256 | Haemophilus<br>parainfluenzae<br>(Bac/Gamma-proteo) (100;0.10) | FQ312002 | Ruminococcus<br>champanellen<br>sis<br>(Bac/Firmicute)<br>(100;0.04) | FP929052 |
| Bifidobacterium<br>animalis<br>(Bac/ActnBac)<br>(100;0.03) | CP009045 | Intestinimonas<br>butyriciproducens<br>(Bac/Firmicute)<br>(100;0.24) | CP011307 | Ruminococcus<br>sp SR1/5<br>(Bac/Firmicute)<br>(100;0.68) | FP929053 |
| Bifidobacterium<br>bifidum<br>(Bac/ActnBac)<br>(100;0.31) | CP010412,<br>CP001840,<br>CP002220,<br>CP001361 | Klebsiella<br>aerogenes<br>(Bac/Gamma-proteo) (91.84;0.01) | FO203355, CP002824 | Ruminococcus<br>torques<br>(Bac/Firmicute)<br>(100;0.97) | FP929055 |
| Bifidobacterium<br>breve<br>(Bac/ActnBac)<br>(97.96;0.01) | CP006715,<br>CP006713 | Klebsiella<br>michiganensis<br>(Bac/Gamma-proteo)<br>(93.88;0.002) | CP004887 | Sphingobacter<br>ium faecium<br>(Bac/CFB_bac)<br>(95.92;0.04) | LK931720 |
| Bifidobacterium<br>dentium<br>(Bac/ActnBac)<br>(85.71;0.01) | AP012326 | Klebsiella<br>pneumoniae<br>(Bac/Gamma-proteo) (88.78;0.01) | CP009208 | Streptococcus<br>mitis<br>(Bac/Firmicute)<br>(100;0.02) | FN568063 |
| Bifidobacterium<br>kashiwanohense<br>(Bac/ActnBac)<br>(100;0.13) | AP012327,<br>CP007456 | Klebsiella variicola<br>(Bac/Gamma-proteo) (90.82;0.01) | CP001891 | Streptococcus<br>parasanguinis<br>(Bac/Firmicute)<br>(100;0.04) | CP002843,<br>CP003122 |
| Bifidobacterium longum<br>(Bac/ActnBac)<br>(100; 0.74) | AP014658,<br>CP002286,<br>CP011964,<br>CP000605,<br>LN824140,<br>AP010890,<br>AP010889,<br>AP010888,<br>CP002010,<br>FP929034,<br>CP006741,<br>CP002794,<br>CP009072 | Lachnoclostridium phytofermentans<br>(Bac/Firmicute)<br>(100;0.09) | CP000885 | Streptococcus pasteurianus<br>(Bac/Firmicute)<br>(100;0.02) | AP012054 |
| Bifidobacterium thermophilum<br>(Bac/ActnBac)<br>(100;0.005) | CP004346 | Lactobacillus acidophilus<br>(Bac/Firmicute)<br>(93.88;0.003) | CP005926 | Streptococcus salivarius<br>(Bac/Firmicute)<br>(100; 0.09) | CP009913,<br>FR873482,<br>CP002888,<br>FR873481 |
| Blautia obeum<br>(Bac/Firmicute)<br>(100;0.51) | FP929054 | Lactobacillus paracasei<br>(Bac/Firmicute)<br>(100;0.01) | AP012541 | Streptococcus sp I-P16<br>(Bac/Firmicute)<br>(100;0.01) | CP006776 |
| butyrate-producing bacterium SM4/1<br>(Bac/Firmicute)<br>(100;0.13) | FP929060 | Lactobacillus rhamnosus<br>(Bac/Firmicute)<br>(92.86;0.01) | CP003094 | Streptococcus suis<br>(Bac/Firmicute)<br>(100;0.02) | CP000837 |
| butyrate-producing bacterium SS3/4<br>(Bac/Firmicute)<br>(100;0.36) | FP929062 | Lactobacillus ruminis<br>(Bac/Firmicute)<br>(100;0.16) | CP003032 | Streptococcus thermophilus<br>(Bac/Firmicute)<br>(100;0.05) | CP000024,<br>CP000419,<br>CP006819 |
| Campylobacter coli (Bac/Delta-Epsilon-proteo)<br>(100;0.01) | CP007180 | Lactococcus lactis<br>(Bac/Firmicute)<br>(96.94;0.01) | CP006766 | Tannerella forsythia<br>(Bac/CFB_bac)<br>(100;0.06) | CP003191 |
| Campylobacter hominis<br>(Bac/Delta-Epsilon-proteo)<br>(97.96;0.003) | CP000776 | Leuconostoc citreum<br>(Bac/Firmicute)<br>(93.88;0.003) | DQ489736 | Treponema succinifaciens<br>(Other Bacteria)<br>(100;0.03) | CP002631 |
| Candidatus Methanomassiliicoccus intestinalis<br>(Arch/Euryar)<br>(34.69;0.01) | CP005934 | Mageeibacillus indolicus<br>(Bac/Firmicute)<br>(100;0.01) | CP001850 | Veillonella parvula<br>(Bac/Firmicute)<br>(100;0.05) | CP001820 |
| Citrobacter<br>freundii<br>(Bac/Gamma-proteo)<br>(84.69;0.02) | CP007557 | Megamonas sp<br>Calf98-2<br>(Bac/Firmicute)<br>(100;0.02) | FP929048 |  |  |
| Clostridioides<br>difficile<br>(Bac/Firmicute)<br>(1.02;2.10) | CP003939,<br>CP010905 | Methanobrevibacter<br>smithii (Arch/Euryar)<br>(39.80;0.07) | CP000678 |  |  |

<sup>1</sup>Percentage of samples this organism is present in.
<sup>2</sup>Average percent relative abundance of this organism.

Several researchers have focused on the reference genes of the gut microbiome rather than organisms [18,39], but organisms have their own clinical significance in treatment. When Yatsunenko et al analyzed 531 healthy samples from Venezuela, rural Malawi and US metropolitan areas and mapped their reads to 126 microbial species, they found Fusobacteria that were not mapped to our list. On the other hand, Spirochaetes, Planctomycetes were not shown in their list [40]. 40 of the organisms reported in their study map to our list at the species level. Unmapped species include organisms such as Actinomyces odontolyticus, Bacteroides capillosus, Bacteroides uniformis and so on. Nishijima et al identified 26 major genera in healthy Japanese [41]. 20 out of 26 genera they listed mapped to our list, the unmapped genera belong to existing GlytFeelingKB families and are Dorea, Dialister, Succinatimonas, Butyrivibrio, Collinsella, and Phascolarctobacterium. Qin et al grouped 66 clusters representing cognate bacterial species for healthy and liver cirrhosis patients [42], and the lowest taxonomy level of cluster in this study is strain. 36 clusters map to GutFeelingKB in the taxonomy levels higher than species and all of them map to existing GutFeelingKB families. It is expected that while other studies will find additional organisms, GutFeelingKB can provide a reference list and abundance information that can provide a starting point for comparative analysis of samples from healthy individuals from around the world and can also help better understand observed differences due to disease and therapy.

#### Organism abundance in individual samples

Many studies have focused on higher taxonomy nodes, providing little species and strain abundance information. Figure 2 shows the abundance of phyla to highlight how our baseline gut microbiome compares to past studies. We provide an abundance sheet with the lowest taxonomy node broken down to the strain level where applicable so that other scientists can use them. Then we calculated the average abundance, standard deviation, maximal and minimal abundance excluding the organisms with the 0% abundance (S10 Table). In terms of average abundance of organisms 4 phyla have abundance above 1%, these are Actinobacteria (1.82± 3 %), Bacteroidetes (73.13 ± 22.16%), Firmicutes (22.2 ± 18.66 %) and Proteobacteria (2.15 ± 10.39%). Bacteroidia (72.97 ± 22.14%) under Bacteroidetes, Actinobacteria (1.67 ± 2.94%) under Actinobacteria, Gammaproteobacteria (2.12 ± 10.38 %) under Proteobacteria, Clostridia (21.35 ± 17.87%) under Firmicutes are the only four classes that have average abundance larger than 1%. Bacteroidaceae (65.58 ± 21.84 %) is the most abundant family, followed by Lachnospiraceae (11.46 ± 11.06%) and Ruminococcaceae (8.38 ± 10.48%). Odoribacteraceae, Rikenellaceae, Bifidobacteriaceae, Enterobacteriaceae and Tannerellaceae are the five other families with abundance above 1%. *Bacteroides* is the most abundant genus in human gut microbiome (65.58 ± 21.84%) with sample SRS016585 having the smallest abundance (0.37%) while SRS013215 has the largest abundance (98.82%). *Bacteroides* includes 9 species and 7 of them have abundance greater than 1%. *Bacteroides dorei* is the most dominant species with a 17.44 ± 8.74% abundance.

**Figure 2.**
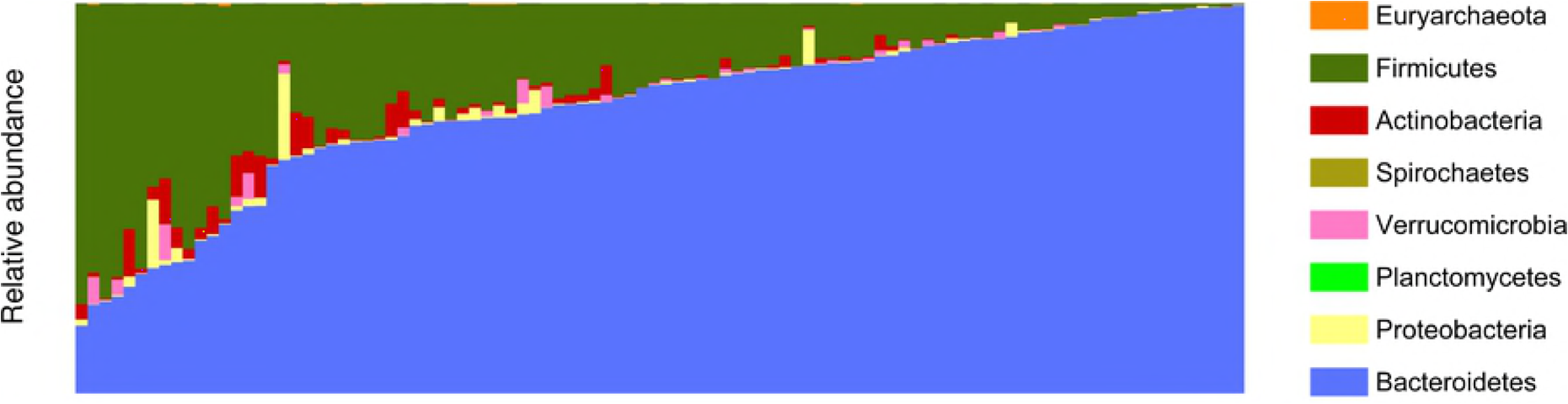
Stacked bar plot of phylogenetic composition of microbiome taxa at the phyla level in fecal (n=98, bottom) samples.

Out of 98 samples analyzed, only 53 samples had archaea. Bacteroidetes, Proteobacteria, Spirochaetes, Actinobacteria, Firmicutes phylum are present in all samples. The abundances of Bacteroidetes are larger than 10% in 97 of 98 samples. *Bacteroides* is present in all the samples with an abundance ranging from 0.37% to 98.82%. Within *Bacteroides*, *Bacteroides fragilis* is present in all the samples. The range of *Bifidobacterium* abundance in all the samples ranges from 0.004% to 12.21%, *Bifidobacterium longum* abundance from 0.003% to 10.30% and *Bifidobacterium bifidum BGN4* strain is present in 96 of 98 samples. A total of 84 out of 109 species are present in all of the samples.

It has been shown that *Bacteroides* is the most abundant genus in Spain, China, Sweden, US, Denmark and France from samples collected from healthy individuals [41]. *Bacteroides* maintain a generally beneficial relationship with the host when retained in the gut but can also be opportunistic pathogens. When they escape the gut environment, they can cause significant pathology, including bacteremia and abscess formation in multiple body sites [43]. Otherwise, they have been shown to have beneficial effects on the host immune system. For example, *Bacteroides fragilis* protects animals from experimental colitis induced by *Helicobacter hepaticus*, a commensal bacterium with pathogenic potential [44]. A large proportion of the *B. fragilis* genome is responsible for carbohydrate metabolism, including the degradation of dietary polysaccharides [45]. *Bifidobacterium* has been reported to be present in almost all healthy human fecal samples. Members of *Bifidobacterium* are among the first microbes to colonize the human gastrointestinal tract and are believed to exert positive health benefits on their host [46]. Many species of *Bifidobacterium* are commonly used as probiotics due to their health promoting properties [47]. Certain *Bifidobacterium longum* strains have been used as probiotics against enterohemorrhagic *Escherichia coli* infection due to the production of acetate, a short chain fatty acid, which upregulates a barrier function of the host gut epithelium [48]. In general, they are able to survive in particular ecological niches due to competitive adaptations and metabolic abilities through colonization of specific appendages. There are 12 strains under *Bifidobacterium longum* species. One strain, BBMN68 has been isolated from the feces of a healthy centenarian living in an area of BaMa, Guangxi, China, known for longevity [49]. Another strain of *Bifidobacterium*, BGN4, was shown to prevent CD4(+) CD45RB (high) T-cell mediated inflammatory bowel disease by inhibition of disordered T cell activation in BGN4-fed mice [50]. Despite the well-established health benefits, the molecular mechanisms responsible for these traits remain to be elucidated.

Some potential pathogenic species appear in our and Yatsunenko et al’s healthy samples [40] like *Streptococcus mitis*, a strain that can cause severe clinical symptoms in cancer patients [51]. Most likely, these organisms are opportunistic pathogens or might be involved in diseases that are not yet fully understood. There are several strains of *Escherichia coli*, but it is generally considered a harmless intestinal inhabitant by being one of the first bacterium to colonize human infants and is a lifelong colonizer of adults [52] although, pathogenic strains of *E. coli* have been implicated in the etiology of health problems such as Crohn’s disease, and ulcerative colitis [53].

#### Contigs from unaligned reads (microbial dark matter)

On average, 50% of the reads from an individual sample could not be aligned to any sequence in Filtered-nt. These unaligned reads were assembled into contigs. Previous work has shown that creation of contigs from unaligned short reads can be used to better understand the actual sequence space represented in metagenomics samples [54]. This “microbial dark matter” remains to be elucidated. Using BLAST on these sequences yielded no significant matches. Given that the average protein-coding density of bacterial genomes is 87% with a typical range of 85–90% [55], and the organisms in our reference list range in size from 1.89 – 6.17 Mb, we chose to look at contigs greater than 10Kb. This value would mean that any single sequence would cover at the very least 0.16% of the organism’s genome, or 0.19% of an organism’s coding region and hence reduce the number of false positive contigs. We were able to assemble unaligned reads into 1,467,129 contigs of which 46,095 have a length greater than 10Kb. After building the contigs, sequences greater than 10,000 nucleotides were all filtered into the same file, and each header was formatted to indicate the sample number, gender, age, and ethnicity of the source. The file is available for download at https://hive.biochemistry.gwu.edu/prd/gfkb//content/unalignedContigsGFKB-v2.0.fasta. These contigs are ideal for new primer design for detailed analysis of the gut microbiome.

### FecalBiome Reporting Template

The effects of the microbiome on health status are growing rapidly and have already spawned FDA approved products at various biotech firms [56]. Some firms have even begun to report microbial composition data to consumers. The formats and parameters for generation of these reports are nonstandardized, limiting their research value. It is necessary to standardize the way that the microbiome is discussed in research and, eventually, in the clinic; the earlier this standardization occurs, the more effective it will be as microbiome science becomes a tool for general research and microbiome medicine moves as close to clinic as genomic medicine. Since there is a need for a cycle moving from bench to bedside and back again, we find value in building a clinical-style report on top of a research tool with the ability to easily cross between the two [57]. This report is also intended to serve as a snapshot of a research project, allowing colleagues and collaborators across labs to share high level information in a rapid manner. Here, we present the FecalBiome Template (Figure 3) -- a general reporting template for microbiome research. It is composed of three domains: Sample, Patient, and Result; these results are drawn from information from a given microbiome sample which is the compared to the contents of the GutFeelingKB. The template was drafted in the spirit of comprehensive metabolic panel (CMP) lab test (https://www.accesalabs.com/downloads/quest-lab-test-sample-report/Comprehensive-Metabolic-Panel-Test-Results.jpg; https://medlineplus.gov/ency/article/003468.htm). It is not uncommon for sample collection, sequencing, and analysis to happen at different locations with different research groups each having a stake in the data produced. In a research setting, the template can serve as a coversheet for shared data, accompanying sequence data to give collaborators a look at their data without having to write scripts for visualizations. While our group works primarily with GI microbiome samples, this report is designed to be generalizable to any human microbiome. Here, we apply the template to the human GI tract, the largest known repository of microbes in the human body.

**Figure 3.**
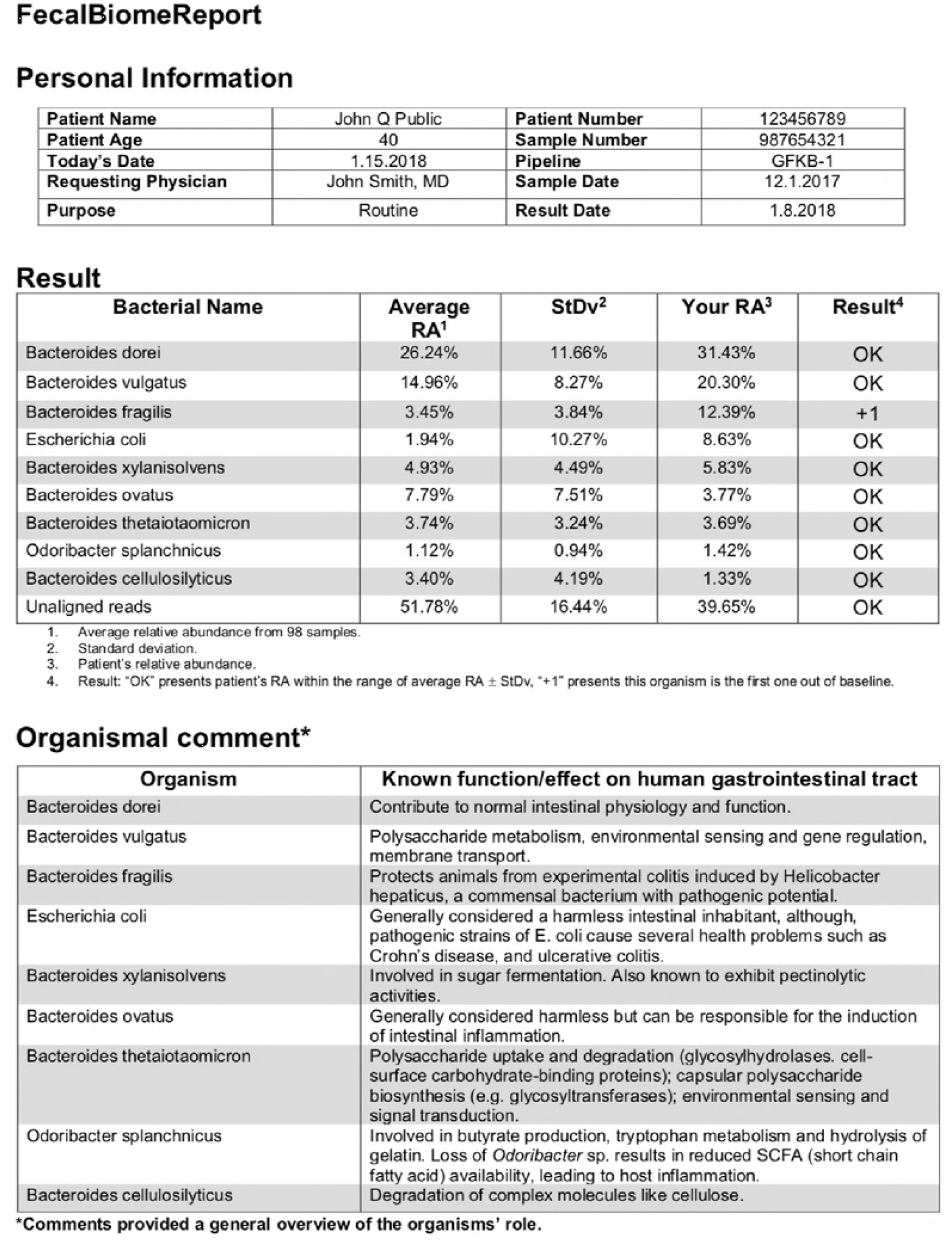
FecalBiome Reporting Template. Personal Information section of the report contains information about the individual who had a sample sequenced, as well as the individual who ordered the sequence. It contains information about the pipeline used for analysis, as well as the sample number for ease of retrieval. Result section contains microbes representing the most abundant organisms which comprise the top 50% of inhabitants. Organismal Comment section includes information from the GutFeelingKB which pertains to the potential function of that organism.

Researchers and clinicians can determine a threshold for the number of organisms reported. Here, we report the organisms comprising the top 50% (sorted based on abundance) of identified microbes from an individual’s sample is included but any threshold of organisms to report in the second and third domains can be set by the user to fit their purposes. Information about abundances, average abundances, as well as information about those microbes from the literature is included on this report.

One of the major outcomes for microbiome science is used in the clinic as any routine test. We intend this report to be the first step in a discussion of standardized reporting of microbiome medicine, bringing the science closer to the clinic (Figure 3). The human GI microbiome is appreciably relevant to human health status. While it is still the early days of microbiome science, it is important to think towards a future where microbiome assays and sequencing are as relevant as routine blood draws and urine samples. As such, we have designed a template for clinical microbiome reporting for physicians and patients. The header of this template was designed to capture relevant information about the test. The two tables which follow the header include the most abundant microbes in a sample, as well as any known physiology and effects of those microbes.

As a test case, we took one sample from the set to determine where it fell relative to the baseline gut microbial population to show the potential clinical application of this technology. For ease of interpretation, the final column in the Result table includes information about whether a given population of microbes falls within the range expected based on the sample space included in GutFeelingKB. The report does not include an explanation for what a particular result means, as it is both premature to tie microbe to phenotype in cases other than infectious disease and any result falls to the purview of the requesting physician. With more information on the role of the microbiome and its constituent microbes, it will become important to be able to compare where a sample from an individual falls within the spectrum of healthy or dysbiotic abundances of microbes.

All relative abundances were calculated for the individual datasets before quantifying the relative min, relative max, mean, median, and standard deviation (Figure 3). These statistics were then transformed into one cohesive report that merged the range, mean, median, and standard deviation. The statistics were further collapsed by family to generate an overall report that models a complete metabolic profile. The top most abundant families (Akkermansiaceae, Bacteroidaceae, Enterobacteriaceae, Rikenellaceae, and Ruminoccocaceae) had a relative max of 8.03, 12.13, 10.99, 6.89, and 6.31 percent of relative abundance, respectively. This is not surprising considering the Rikenellaceae family is indicative of good gastrointestinal health [58]. Akkermansiaceae is linked to lower rates of obesity and associated metabolic disorders [59]. Bacteroidaceae and Enterobacteriaceae can be linked to acute infective processes but are otherwise symbionts [60,61], and Ruminococcaceae is known to break down complex carbohydrates especially in people with carb heavy diets [62]. FecalBiome and the underlying GutFeelingKB can have high value to clinicians who hope to assess the gut microbial status of their patients. The goal of the database and report is to connect lab results with outcomes. At present, most microbiome diseases are those of severe dysbioses caused by a kind of potentially pathogenic bacteria – the canonical infectious pathogens such as *Helicobacter pylori*, *Vibrio cholerae* and others. By determining what species or strain correlate with good or bad outcomes, we could aid clinicians in developing strategies for valuable evidence-based treatments.

### Dietary data and nutrient correlative analysis

MaAsLin is a multivariate statistical framework that identifies associations between clinical metadata and microbial community abundance [36]. The HMP samples did not have specific dietary data (participants only categorized their type of diet: Carnivore, Vegan, Vegetarian etc.), and thus, this analysis was limited to the samples collected at GW. Over 100 features were obtained for each participant from the NDSR program and added this to the abundance sheets, along with the anthropomorphic measurements (height, weight, waist circumference) that were taken.

In comparing bacterial species to nutrient data, several interesting patterns were observed. *Bifidobacterium* was positively correlated with dietary protein intake (Figure 4a), specifically vegetable protein, as well as dietary fiber, specifically soluble fiber, present in vegetables such as broccoli, brussel sprouts, beans, peas, asparagus and beans, which also contain vegetable protein. *Akkermansia* (figure 4b) was shown to be positively associated with saturated fat intakes and is negatively correlated with total polyunsaturated fatty acids (PUFA). Not surprisingly, it was also positively correlated with linoleic acid, as this particular omega-6 PUFA is found abundantly in oils (e.g. soybean oil, vegetable oil) used in processed food. *Bacteriodes ovatus* was positively correlated with daily calorie intake (Figure 4c), as well as body weight (Figure 4d), and waist circumference. The table of results (see supplementary table S11 Table) demonstrates the range of correlation for features that have been measured.

**Figure 4.**
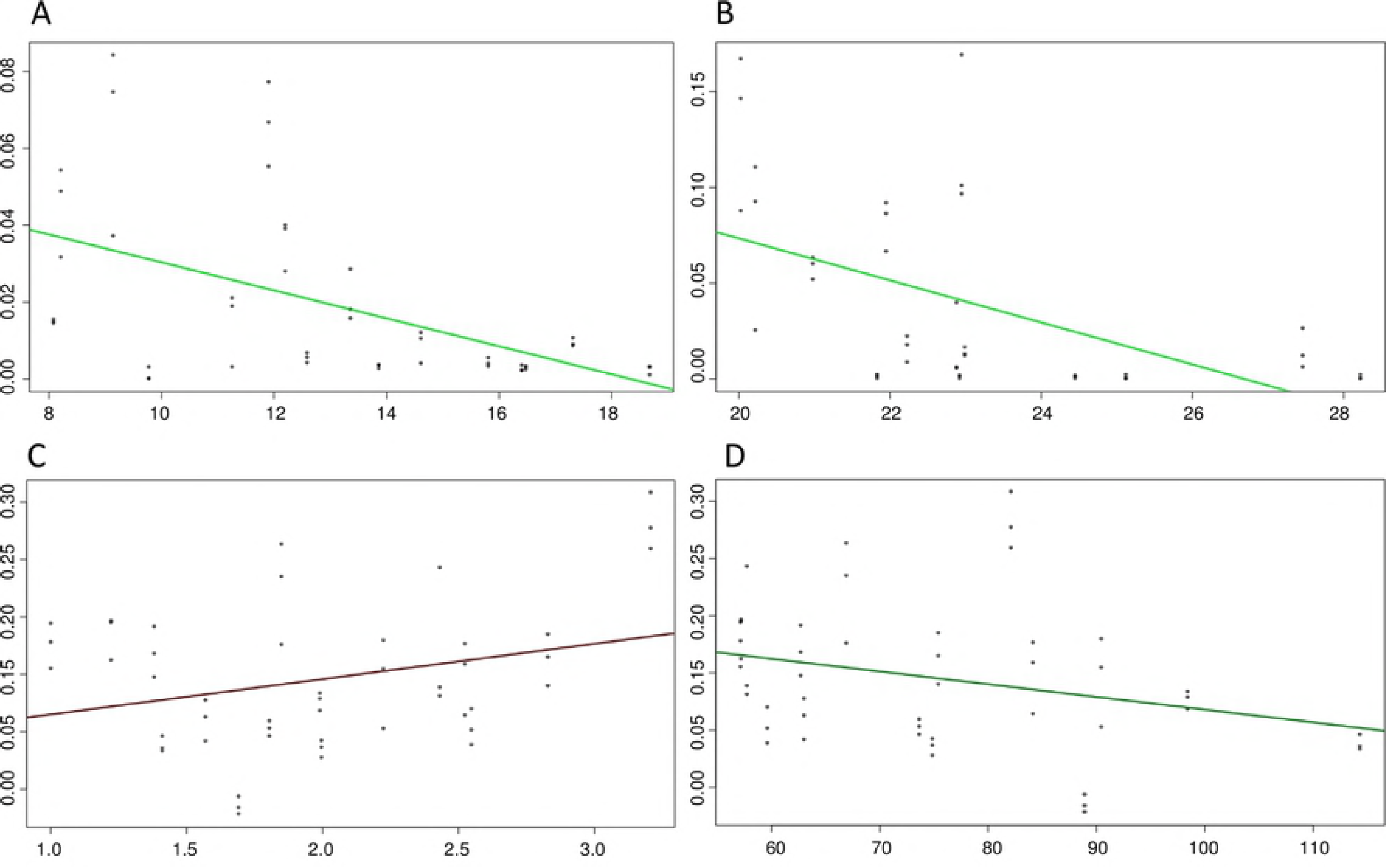
Correlation between bacterial organism and nutrient data. (A) *Bifidobacterium* is positively correlated with dietary protein intake, specifically vegetable protein, present in vegetables such as broccoli, brussel sprouts, beans, peas, asparagus and beans. (B) *Akkermansia* is positively associated with body mass index (BMI). (C) *Bacteriodes ovatus* is positively correlated with daily calorie intake. (D) *Bacteriodes ovatus* is negatively correlated with daily body weight.

Cosine Similarity Coefficient analysis (see supplementary table S12 Table) identified correlation for features and organisms with the observations similar to MaAslin. For example, characteristics such as fat intake and BMI correlate with members of *Akkermansia*. Similarly, the impact of Vitamin A or beta carotenes has positive inductive correlation across all the Bifidobacterium (Figure 5).

**Figure 5.**
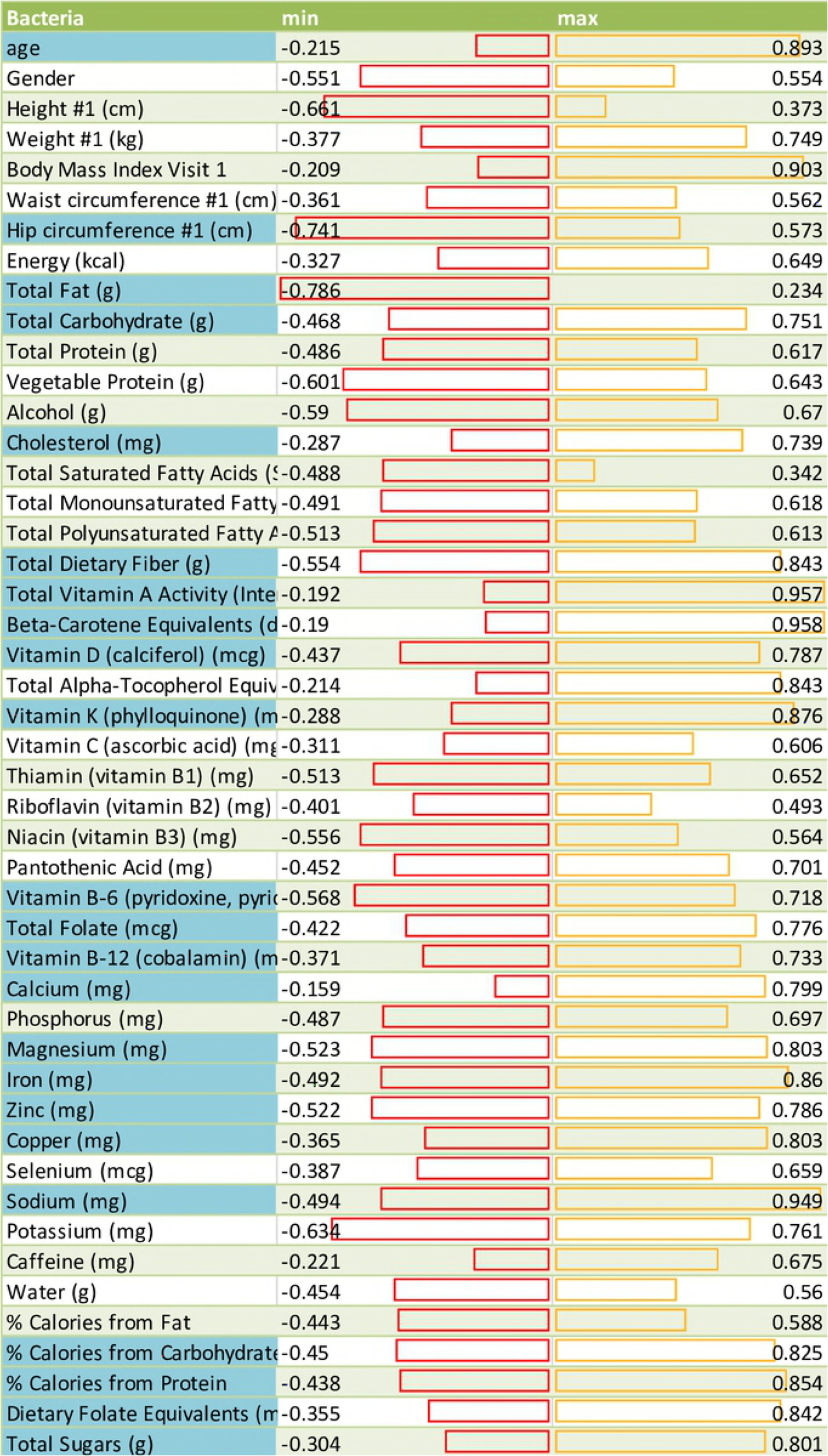
The range of correlation for all features that have been measured for each of the GW samples. Each line is a graph of the min and max values using a Cosine Similarity coefficient correlation. A positive value means strong correlation, and a negative value means strong anticorrelation, whereas zero means absolutely no correlation. Given the size of sample pool of 16 we have taken 0.7 as the marginal threshold for evidence of some degree of correlation. Each feature that had a correlation with any organism is highlighted in blue. For example, some characteristics such as Fat intake have anticorrelation with members of *Campulobacter jejuni* and Eubacterium family.

As microbiome science moves closer to the clinic, it will be imperative both to have tools for analysis and the quick understanding of a microbial population. We envision our database and pathway analyses as the foundation for this clinical reporting. While each organism in an entire microbiome sample isn’t immediately actionable, it does allow for both the close tracking of microbial modulation and the better understanding of how the microbiome tracks with health states and therapy. This will be further applicable as evidence based medicine approaches microbiome science, and microbiome science becomes as important to clinical treatment as genomic medicine. Preliminary microbiome analyses are increasingly yielding interesting results in complex diseases such as cancer. For example, in colorectal cancer patients, carcinoma-enriched bacteria, *B. massiliensis*, *B. dorei*, *B. vulgates*, *Parabacteroides merdae*, *A. finegoldii* and *B. wadsworthia*, positively correlated with red meat consumption and negatively correlated with fruit and vegetables consummation [63]. It is expected that as the number and size of these studies increase, the need for baseline human gut microbial profile in healthy people and standard reporting template will become essential.

### Conclusion

The workflow described in this study involves a sub-sampling-based method followed by comprehensive mapping of all of the reads to accurately determine the abundance of microorganisms. The workflow provides a comprehensive snapshot of the microbial abundance and can easily be used with any state-of-the-art NGS read mapping and assembly algorithm. The list of baseline organisms identified in the normal human gut has clinical applicability as microbiome research moves closer to the bedside. The methods, tools and data from this project can also be used by regulatory scientists to evaluate workflows related to fecal transplant.

In addition to the workflow, we have laid the foundation for an expansive and modular database which will aggregate all publicly available data as well as the data from contributors to push towards an understanding the baseline human microbiome. This database will serve as a common control in studies of dysbiosis and microbiome associated common disease and cancer. Finally, the user-friendly format through FecalBiome report, which contains absolute and relative abundance information about a given sample compared to an average across the entire database, scientists, clinicians, and eventually patients can get an easy to understand overview of gut microbiome. Separately, we see a significant impact of this technology on regulatory science in the future. Finally, as a tool and library, GutFeelingKB will allow for rapid assessment of the content of human GI replacement products and, ideally, allow for more expedient review of products. Future studies to advance evidence-based microbiome medicine should be conducted where potential patients identify which outcomes such as depression, bloating, frequency of common colds, etc., are most important via a focus group or survey. Those outcomes will become endpoints in clinical trials or observational studies that demonstrate the effects of various bacteria on the human gut. This type of methodology would tie raw numbers to health states that are meaningful for the general population, ensuring that data gathered are relevant to the patient, and therefore the clinician. This could bring a new, patient-centric perspective to microbiome data and allow for a greater scope of health data to sit atop metagenomic sequence data. These outcomes/endpoints would become a “toolkit” for other researchers who are interested in the gut. If everyone uses the same set of clinically relevant endpoints, research will be easily comparable across studies and meta-analysis becomes interoperable.

## ACKNOWLEDGEMENTS

We would like to thank the following people for providing valuable comments: H Zhang and Y Hu. We would also like to thank all the students have contributed to the GutFeelingKB organism descriptions over the years. This project was supported in part by funds from National Science Foundation (NSF) (award number: 1546491 to RM) and the McCormick Genomic and Proteomic Center (MGPC) at the George Washington University.

## Supplementary files legends

S1 Table. Anthropomorphic measurements of GW and HMP samples. 1.) GW anthropomorphic measurements and the associated value. 2.) HMP anthropomorphic measurements and the associated value. 3.) Selection criterions

S2 Table. Nutritional features and the associated values of GW and HMP samples. 1.) GW 100 nutritional features and the associated values from NDSR results of GW samples. 2.) 100 nutritional features and the associated values of HMP samples.

S3 Fig. Quality assurance of one sample. (A) Summary statistics for the read file. (B) ACGT Count: A pie chart displaying the number and percentage of bases present in a read file. (C) Lengthwise Position Count: Displays the number of bases versus position in the read files. (D) Quality Position Count: The average quality score of a position in the reads of a file. (E) Average Quality Per Base: A histogram of the quality score of each base pair. (F) Length Count: A plot of the read length against the number of reads in the sample. (G) Quality Length Count: Shows the average quality score of a read of a given length.

S4 Fig. HIVE-MultiQC output figures. (A) The average quality score for each base shown by sample file. The consistently high-quality score for the forward strand files indicates acceptable sequences for analysis. (B) The relative abundance of each base in each read file. (C) The average quality score for the entire data set, shown by position in the read, is the blue line. The greyed area represents one standard deviation above and below the average.

S5 Table. GW read files information. List of 96 reads file information from 48 GW samples.

S6 Fig. Enterotypes of GW and HMP samples.

S7 Table. Mapping information of the organisms in GutFeelingKB. Organisms shown in GutFeelingKB are present by UniProt IDs, all the UniProt IDs have been mapped to NCBI. This table lists the same organism’s information through different databases like UniProt, NCBI Assembly, NCBI Taxonomy, NCBI Nucleotide and so on.

S8 Table. Pathogens table. List of well-known gut pathogens that can be misidentified through metagenomics.

S9 Table. Blacklist of Filtered-nt. All the removed taxonomy IDs all shown in this table.

S10 Table. Abundance table. Abundance table are presented by 7 tables in deferent taxonomy level, including phylum, family, class, order, genus, species and strain abundance tables. Average abundance, standard deviation, maximal and minimal abundance are provided excluding the organisms with the 0% abundance.

S11 Table. Associations between clinical metadata and microbial community abundance.

S12 Table. Cosine similarity coefficient of correlation. This table demonstrates the range of correlation for features that have been measured, and the organisms that have been detected.

